# Phenotype-dependent schooling behaviour across hierarchical levels in three-spined sticklebacks

**DOI:** 10.64898/2026.09.17.752332

**Authors:** Aparajitha Ramesh, Eize Stamhuis, Marion Nicolaus

## Abstract

Schooling is a widespread behaviour among pelagic fish that reduces locomotion costs of migration and predation risk. It is characterised by coordinated movements, speed and orientation of group members. Although schooling can vary both between populations and among individuals of the same population, these levels of variation are seldom considered together. Here, we investigated whether schooling tendencies in three-spined sticklebacks (*Gasterosteus aculeatus*) vary between populations and among individuals within-populations depending on their phenotype. We first compared the schooling behaviour of anadromous (‘migrants’) and recently landlocked (‘residents’) populations and tested whether individual sociability explained within-population variation in schooling. We then studied whether schooling behaviour is modulated when migrants and residents are in mixed groups. Migrant groups displayed better schooling performance than residents with higher alignment. Within migrant groups, less social individuals displayed more leadership, were better aligned, swam closer to others and fatigued less than more social fish. These patterns were absent in residents. When the two populations were mixed, overall schooling performance declined, particularly in migrants, with schools exhibiting reduced compaction and alignment, resembling resident groups. Thus, individuals did not modulate their schooling behaviour to the group social composition and phenotype-dependent schooling patterns were disrupted in migrants when mixed. These findings suggest that migration-related behaviours can disappear quickly post-isolation. They also highlight the importance of individual sociability in self-organisation and shaping collective behaviour, especially in populations that retain migratory behaviour. Considering behavioural variation across multiple hierarchical levels is thus key to understanding complex social systems.

**Significance statement:** Many animals exhibit complex social behaviours that can benefit group members. In fish, schooling behaviour enables to migrate with reduced costs of locomotion and predation risk. Yet, the benefits of schooling and schooling behaviour itself are shaped by the environmental conditions, and thus are strongly influenced by human disturbance. We conducted a study in three-spined sticklebacks to test whether man-made barriers that prohibit some fish populations to undergo their seasonal migration (‘residents’) altered schooling behaviour compared to their ‘migrant’ anadromous ancestors. Migrants were indeed better at schooling than residents, and their schooling patterns were shaped by individual variation in sociability. Residents did not improve their schooling behaviour when mixed with migrants. This suggests that ∼ 50 years of isolation are sufficient to losing migration-related behaviour and that river reconnection will likely promote the emergence of partial migration in this system.

## Introduction

Among all vertebrates, fish provide some of the most spectacular examples of social aggregation with their ability to form schools that can exceed several million individuals in species like the Atlantic herring (*Clupea harengus*). Unlike shoals, which are defined exclusively by social attraction, schools are characterised by individuals maintaining common orientation and synchronised speed and movements (Kasumyan and Pavlov 2023a). Schooling provides advantages for the individuals, such as reduced predation risk, increased foraging opportunities or reduced costs of locomotion (Kasumyan and Pavlov 2023b, Zhang and Lauder 2024, Zhang et al. 2024). For example, in grey mullets (*Liz aurata*), individuals in a school experience up to 20% reduction of energetic costs compared to solitary swimming (Marras et al. 2015). Accordingly, schooling is often observed in migratory and pelagic species that must travel between areas with few shelters, while facing high fitness costs related to energy expenditure and predation risk (Kasumyan and Pavlov 2023a, 2023d).

Schooling behaviour has been shown to have a genetic component in some species, implying that it can evolve under natural and artificial selection (Kotrschal et al. 2020, Corral-Lopez et al. 2023). Consequently, when a fish population can no longer migrate and becomes spatially restricted, e.g. due to physical isolation in small water bodies, the net benefits of schooling could decrease and, ultimately, non-migratory populations could exhibit lower schooling tendency and differentiate genetically from their anadromous ancestors in these traits. For example, in three-spined sticklebacks (*Gasterosteus aculeatus*), individuals from benthic lake habitats have evolved a reduced schooling tendency compared to marine conspecifics from pelagic open habitats (Wark et al. 2011, Greenwood et al. 2016). Similarly, cave and surface forms of Mexican tetra (*Astyanax mexicanus*) have diverged behaviourally and genetically, with cave populations no longer exhibiting the tendency to school (Kowalko et al. 2013).

Schooling behaviour is an emergent characteristic of fish groups, and hence is typically studied at the species or population level. However, schooling actually arises from the behaviour of individual fish that possess their own phenotypic attributes, including differences in behavioural tendencies (or personality) (Herbert-Read et al. 2013). Therefore, it has been argued that these individual differences may drive the behaviour and functioning of groups and should be considered as a key aspect of collective animal behaviour (Viscido et al. 2004, Jolles et al. 2020). Empirical evidence shows that individual personality can indeed predict individual schooling ability (Jolles et al. 2017). For example, several studies found that bolder and less social individuals tend to lead schools more often and maintain larger inter-individual distances than shyer and more social individuals (Ward et al. 2004, Jolles et al. 2017, Bevan et al. 2018). In grey mullets, schooling individuals were also found to differ consistently in their escape latency under predation threat (Marras and Domenici 2013). Modelling studies have further shown that the composition of the school matters for its performance: typically, the more dissimilar the individuals (e.g. in terms of behaviour or size), the slower, less compact, aligned or coordinated the school is (Kunz and Hemelrijk 2003, Del Mar Delgado et al. 2018). This can be explained in part by individual heterogeneity enhancing assortment by phenotype within schools (Krause et al. 2000). These school properties may, in return, impact the survival or performance of their members (Dyer et al. 2009, Ioannou et al. 2012).

Despite the growing awareness that within-species variation in schooling behaviour can originate from population divergence and the phenotypic composition of individuals comprising the groups, to our knowledge, these two levels of variation have not been considered simultaneously in the same study. Therefore, this study aims to test the extent to which variation in schooling properties can be explained by populations and individual phenotypic variation within populations. To that end, we compared the schooling behaviour of three-spined stickleback groups varying in their social composition. Our study system in the Netherlands consists of ancestral anadromous populations (‘migrants’) and replicates of forced resident populations that can no longer access the sea due to anthropogenic barriers built 50-60 generations ago (‘residents’). Previous studies have shown that these populations, while harbouring large personality variation, diverged behaviourally, with resident fish appearing to be on the way to losing migration-related behaviours (Ramesh et al. 2021, 2022). In addition, when the populations were mixed in large mesocosms, no social modulation was detected (i.e. individual fish retain their behavioural tendency) (Gismann et al. 2024).

In this study, we conducted an experiment in which we quantified schooling properties (body orientation, inter-individual distance, position within school and fatigue behaviour) of groups and individuals within groups. Across trials, we manipulated the group composition in terms of population of origin (pure groups of migrants or residents or mixed groups) and personality (activity, aggression, boldness, and sociality). Between the pure resident and migrant groups, we predicted that migrant groups would be more aligned, i.e. swimming more parallel to each other’s (Wark et al. 2011, Di-Poi et al. 2014, Greenwood et al. 2016), more compact, i.e. swimming closer to each other’s (Jolles et al. 2017) and exhibit higher swimming endurance (Greenwood et al. 2013) than resident groups. Within groups and independently of the population of origin, we expected that ‘proactive’ individuals, characterized by relatively higher levels of activity, aggression, boldness and/or reduced social tendency, would take more risks and occupy more often the leading position of the school compared to ‘reactive’ individuals with opposite behavioural characteristics (Ward et al. 2004, Jolles et al. 2017, Bevan et al. 2018). Proactive fish may exhibit relatively larger inter-individual distances due to their faster swimming speed (Jolles et al. 2017) and be more aligned (Tang and Fu 2020), but fatigue more, as individuals swimming at the front may experience higher energetic costs than fish swimming behind their neighbours (Marras et al. 2015). When migrant and residents are mixed, group schooling performance may deteriorate because of resident fish being unable to keep up with the swimming performance of migrants (Gismann et al. 2024), thereby increasing individual heterogeneity within groups (Kunz and Hemelrijk 2003, Del Mar Delgado et al. 2018). In these mixed groups, migrant individuals are expected to take the lead more often compared to residents due to their higher swimming ability. Lastly, ‘reactive’ individuals within mixed groups may show more plasticity than ‘proactive’ conspecifics (Jolles et al. 2019); i.e., they are expected to exhibit a higher degree of social modulation and to adjust their schooling behaviour according to the group composition.

## Material and Methods

### Study animals

The F1 sticklebacks of the migrant and resident populations used in the current study were originally bred for a previous study (see details in Ramesh *et al*., 2021). In brief, F0 anadromous migrants were captured at the mouth of the Westerwoldse Aa River (“NSTZ”; 53°13’54.49” N, 7°12’30.99” E), whereas residents were sampled from two land-locked polders (“LL-A”: 53°17’56.14” N, 7°02’1.28” E; “LL-B”: 53°17’16.52” N, 7°02’26.46” E). We have previously showed that the resident populations did not differ in behaviour (Ramesh *et al*., 2022) and therefore these individuals have been pooled. F1 juveniles were obtained using a partial factorial breeding design with in-vitro fertilisation (following Arnott and Barber 2000). This design yielded pure migrant (M), pure resident (R), and hybrid (H) crosses (H not being used in the current study). F1 offspring were raised under common-garden conditions without parental care. As the fry developed, food and housing conditions were adjusted to body size: larvae were fed frozen cyclops, freshly hatched *Artemia nauplii*, and GEMMA Micro 75, later transitioning to ad libitum brine shrimp and bloodworms (3F Frozen Fish Food bv.). Once juveniles reached ∼2 cm, fish from different families were regrouped into mixed-family tanks of ten, keeping cross types separate.

Fish were maintained in a flow-through system at ∼16°C under a 16:8 h light:dark photoperiod during growth. When individuals reached ∼4 cm, each received a unique identification tag for the previous study (see Ramesh *et al*. 2021). To induce a non-breeding state, autumn conditions were initiated at ∼12–13 months of age by reducing the light period to 12:12 h (L:D) and lowering temperatures to ∼13–14°C. These conditions were maintained until the end of the experimental period, when the fish were ∼15-16 months old in 2020.

### Personality assays

Individual personality scores used in the present study were obtained from standardised laboratory assays conducted previously on the same fish 5 months prior to schooling assays (Ramesh *et al*., 2021). Briefly, individuals were tested twice, at least four weeks apart, in a series of four behavioural assays: activity, exploration, sociability, and boldness. For this study, we focussed on sociability as a key personality trait, due to its direct relevance to schooling behaviour. Sociability (referred to as ‘shoaling’ in the previous study) was measured as the proportion of time the focal fish spent near a conspecific shoal compartment to the total test time in a choice test with a group vs two distractor fish. Sociability was found to be moderately but significantly repeatable in both populations (R = 0.31, Ramesh *et al*., 2021), consistent with the expected range for behavioural traits (Bell *et al.,* 2009). F1 migrants were found to exhibit significantly higher sociability than F1 residents. This indicates that behavioural population difference in sociability is likely underpinned by genetic differentiation, although social effects cannot be excluded. In contrast, other personality traits showed weaker or less consistent patterns between populations (Ramesh *et al*., 2021). Therefore, sociability provided the most reliable and biologically meaningful measure for this study. The two rounds of sociability scores were averaged for each individual.

### Schooling assay - Setup

To quantify variation in schooling behaviour among groups of fish, we tested them in a custom-built flow chamber (Fig. 1). The controlled flow provided a biologically relevant and standardised swimming environment that mimics the directional swimming conditions experienced during migration, while allowing us to quantify schooling behaviours. The setup consisted of a rectangular glass aquarium containing an open Plexiglas tube connected to PVC tubing and a motor-driven pump that circulated water in a closed loop. A laminar flow was generated inside the tube and maintained by honeycomb flow-straighteners positioned at the upstream entrance. A metal grid prevented fish from swimming further upstream. The swimming arena within the tube measured 11.9 cm in diameter and 40 cm in length. The chamber was filmed simultaneously from the side and from a top-down mirror using a GoPro HERO8 (GoPro Inc.). The camera was positioned approximately 20 cm from the aquarium wall (plus 6.8 cm to the Plexiglas tube), and its placement was standardised across trials. A single overhead light source illuminated the arena while all other light directions were blocked to avoid shadows. Blinds surrounding the chamber minimised disturbance.

**Figure 1:**
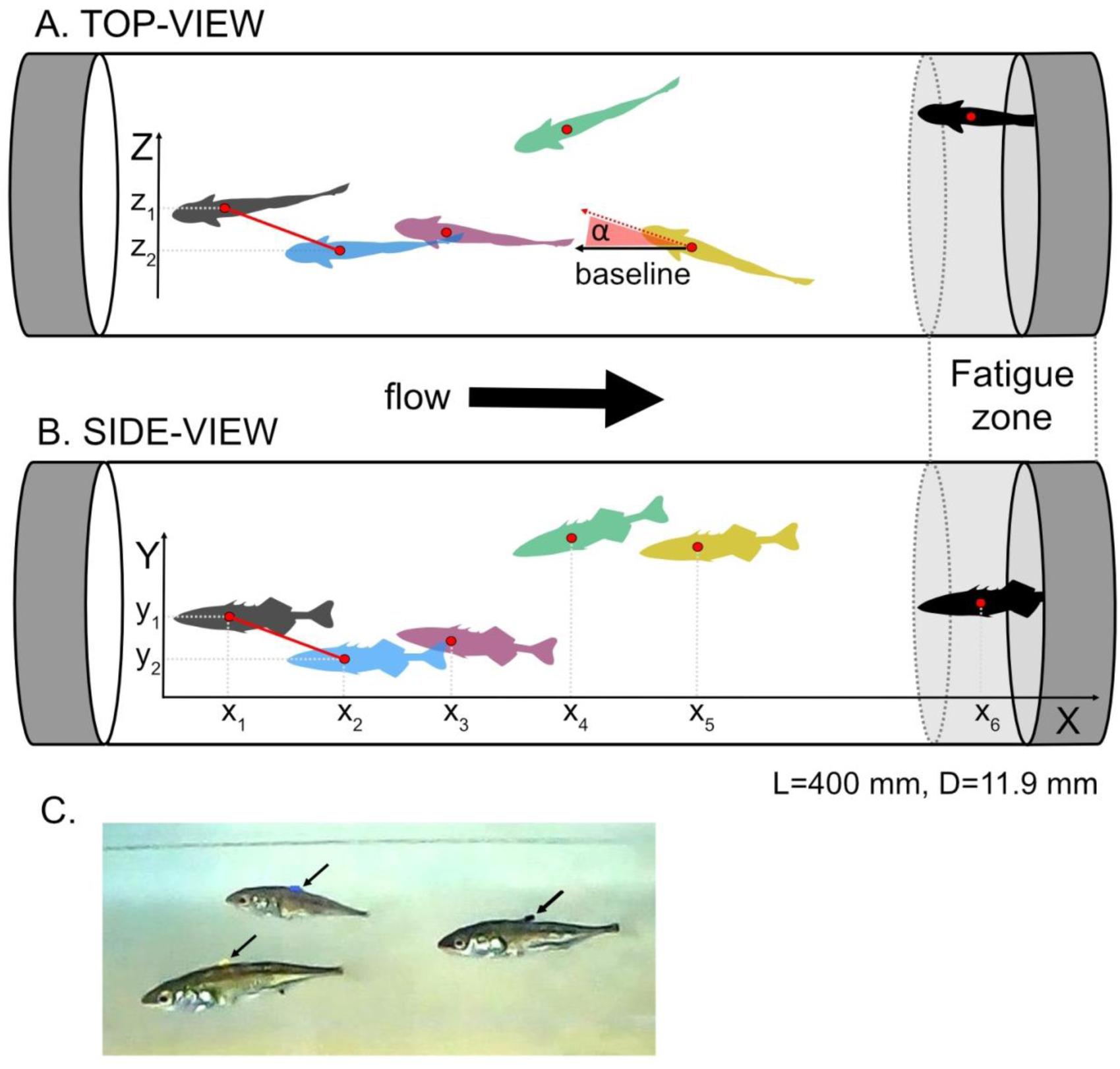
Schematic representation of the top-view (A) and the side-view (B) of the flow chamber with a group of six schooling fish. Length and diameter of the chamber are indicated. Each fish was tagged with a unique colour on its dorsal spine (C). In this video snapshot, colour-tagged yellow, blue and black sticklebacks are seen (left to right).

### Schooling assay - Group composition

Each trial consisted of six randomly selected fish representing different group compositions: Pure groups of migrants or residents, and mixed groups of either majority migrant (4 migrant and 2 resident fish) or majority resident (4 resident and 2 migrant fish). The six fish were chosen from three different home tanks such that a pair of fish originated from each home tank, ensuring familiarity within pairs but also ensuring a constant proportion of unfamiliar fish in both pure and mixed treatments.

Pure (migrant or resident) and Mixed (majority migrant or majority resident) groups were tested in 10 independent trials over a span of 12 days. Total sample size is thus 40 trials (10 pure migrant, 10 pure resident, 10 majority migrant and 10 majority resident). In total, 64 individual fish were used (32 M and 32 R). Most fish were used four times (mean ± SD = 4 ± 1.6, range = 1 to 6 repeats) with a minimum of 3 days between the trials. Because we aimed to first characterised variance in schooling behaviour within and between populations and within and between individuals of the same population before examining plastic adjustments to a change in the social environment, mixed groups were always tested after the pure groups.

### Schooling assay - Testing procedure

To reduce handling stress, fish underwent two acclimation phases. First, the six focal fish were transferred from their home tanks to a similar tank containing a 1:1 mixture of home-tank and tap water for a 10-minute acclimation period. Each individual was then fitted with a unique coloured identification tag in a non-invasive procedure (electric wire insulator placed over the second dorsal spine; Fig. 1C) and then introduced into the flow chamber (water temperature 13.0–15.3°C, matching home-tank conditions). A flow speed of ∼4.0 cm/s was initiated, followed by a 20-min acclimation period. After acclimation, recordings were started (1440p, 60 fps), and the flow speed was increased to ∼6.2 cm/s. Each trial lasted 11 min. Afterwards, fish were placed in the mixed-water holding tank for a further 20- min recovery period before returning to their home tanks.

### Data extraction from videos

For each trial, eleven frames were extracted one minute apart. The first four frames of each recording and the final frame were discarded due to disturbance during setup and shutdown, leaving seven frames per trial for analysis. Total sample size was thus 1680 observations (4 treatments × 40 trials × 7 frames × 6 fish). All measurements and landmark extraction (see below) were performed in ImageJ (NIH and LOCI, University of Wisconsin).

3D positions were estimated from the side and top-view 2D images; each frame was scaled to real-world dimensions. To obtain fish positions, each individual was identified by its unique colour tag and manually located in both views. The X and Y coordinates of each fish were extracted from the side view (Fig. 1B), and the Z coordinate from the top-view (Fig. 1A). Body orientation was quantified from top-view images by measuring the angle of each fish relative to a baseline aligned with the flow axis (angle α in Fig. 1A). Based on these positions and angles, we derived the following measures at the individual and group levels.

#### Individual level

1. *Mean inter-individual distance* was computed as the mean pairwise distance (nearest to fifth nearest neighbour) from the centre of standard length for each fish with respect to the others (Fig. 1A & B). Thus, each focal fish was assigned five inter-individual distances, which were then averaged to give a mean inter-individual distance per individual per frame. Higher values indicate that the focal fish was farther from the group, either because it moved away from others or because the other fish moved away from it.
2. *Individual alignment* was expressed as the deviation of each fish’s angle from the mean school orientation within the frame (individual deviation from mean α, Fig. 1A). Therefore, 0 indicates perfect alignment, and increasing (absolute) deviation from 0 indicates increasing misalignment with the group’s mean. Because individuals are tested with a flow, better alignment may mean better orientation and better polarisation with other group members.
3. *Individual probability to fatigue* was calculated as a binary variable (fatigued / not fatigued) in each frame. A fish was considered fatigued if it reached the far right of the flow chamber (one fish length from the back, i.e. 4.5 cm) (Fig. 1B).
4. *Rank* of each fish within the school was calculated using the position of each fish on the X coordinate, with smaller X coordinates indicating the front position and larger X values the back of the school (X1, X2, etc. on Fig. 1B).

#### Group level

1. *Group compaction* was calculated as the distance between the first and last fish in a school (X6-X1, Fig. 1B)
2. *Group alignment* was calculated as the standard deviation of all individual alignment values (or α) in the school within the frame.
3. *The proportion of fatigued fish* was calculated as the ratio of fatigued fish to the total fish in the school.

### Statistical analysis

#### Individual level

At the individual level, we aimed to quantify variation in schooling traits across multiple levels and sources of variation. First, we tested whether individual schooling behaviour was phenotype-dependent (sociability) within pure groups of migrants and residents. We analysed variation in *mean inter-individual distance*, *individual alignment*, and *rank* using Linear Mixed Models (LMMs) with a Gaussian error distribution and variation in the *probability to fatigue* using a binomial Generalised Linear Mixed Model (GLMM). In all models, individual sociability (relative to the group mean), mean group sociability, population (migrant vs. resident) and the interaction (population × sociability) were included as fixed effects and Fish ID, Group ID (= Trial ID) and Trial frame ID as random effects to account for repeated measurements and interdependence of data. Including all these variables in the same models was possible because despite mean population difference in sociality, substantial individual variation remains within population and some overlap exists between residents’ and migrants’ values (Ramesh *et al*., 2021). Since individuals were assigned in a group based on their origin and not their level of sociality, variation exists between group means, even within the same population. This allowed us to separate within- and between-group effects of sociability.

Next, we tested whether migrants and residents modulate their behaviour when schooling in pure vs mixed groups. To this end, individual schooling traits were used as response variables, with population (migrant vs. resident), treatment (pure vs. mixed), and their interaction included as fixed effects. Fish ID, Group ID (= Trial ID), and Trial frame ID were included as random effects. Mean inter-individual distance, individual alignment, and rank were analysed using LMMs with a Gaussian error distribution, while probability of fatigue was analysed using a GLMM with a binomial error distribution and logit link function.

Finally, to test whether specific personality types (more social fish) and/or a specific population were more plastic and adjust to changes in the social environment, we tested different individual schooling traits with relative individual sociability (centred on the group mean), mean group sociability, population origin (migrant vs. resident), treatment (pure vs. mixed), and a three-way interaction between population, sociability, and treatment included as fixed effects, and Fish ID, Group ID (= Trial ID), and Trial Frame ID included as random effects. This will enable us to disentangle how individuals differing in sociability in migrant or resident groups change when their social environment changes with respect to schooling traits. The main difference with the previous models described above lays in the fact that the focus is placed on the behavioural changes between homogenous (same origin, pure migrant and pure resident merged) and heterogeneous (mixed groups) social environments, a relevant contrast given the ongoing effort to reconnect the populations in the wild.

Note that influence of trial order on schooling metrics was tested in all models but had no significant effect (not shown). Therefore, it was removed to keep the models most parsimonious

#### Group level

At the group level, we aimed to test whether groups of pure migrants and residents differed in their schooling behaviour and whether they also differed from a mixed group comprising both migrants and residents. To this end, the different schooling traits, including *group compaction* and *mean group alignment,* were modelled as response variables with treatment (migrant, resident or mixed) included as fixed effects and Group ID (= Trial ID) included as a random effect in LMMs with a Gaussian error distribution. Variation in the *proportion of fatigued fish* was further analysed using a binomial GLMM with a logit link function and the same fixed and random effects structure. We initially analysed the majority-migrant and majority-resident treatments in mixed groups separately, but decided to pool the data into mixed groups on finding no difference between the two mixed groups.

All LMMs/GLMMs were constructed in R v.4.5.0 (R Core Team 2025) using the “lmer” function of the “lme4” package (Bates et al. 2015). Normality of all dependent variables was checked prior to the analyses. Binomial models were tested for overdispersion using the “Dharma” package (Florian et al. 2017) and they were not overdispersed. The statistical significance of fixed effects was assessed based on the 95% confidence interval (CI): an effect was considered significant when its 95% CI did not include zero. Adjusted repeatabilities (after accounting for fixed effects), and their confidence intervals were calculated using “rpt” function with 1000 bootstraps in “rptR” package (Stoffel et al. 2017). Figures were produced using “ggplot2” package (Wickham 2016).

## Results

### Personality and individual schooling behaviours in pure groups

Within pure groups, we found that the individual social tendency predicted variation in individual schooling behaviours and that the strength of these relationships was population-specific. In migrants, relatively less social individuals occupied the front position of the school more often, at a larger distance from their conspecifics and in a more aligned fashion compared to relatively more social individuals (Table 1, Fig. 2). These asocial individuals also tended to be more likely to fatigue (Table 1, Fig. 2). Mentioned trends were significant when models were run separately for migrants (not shown). In contrast, within resident groups these patterns disappeared, i.e. individual social tendencies did not predict schooling behaviours (see two-way interactions in Table 1, Fig. 2), even when models were run separately for residents (not shown).

**Figure 2:**
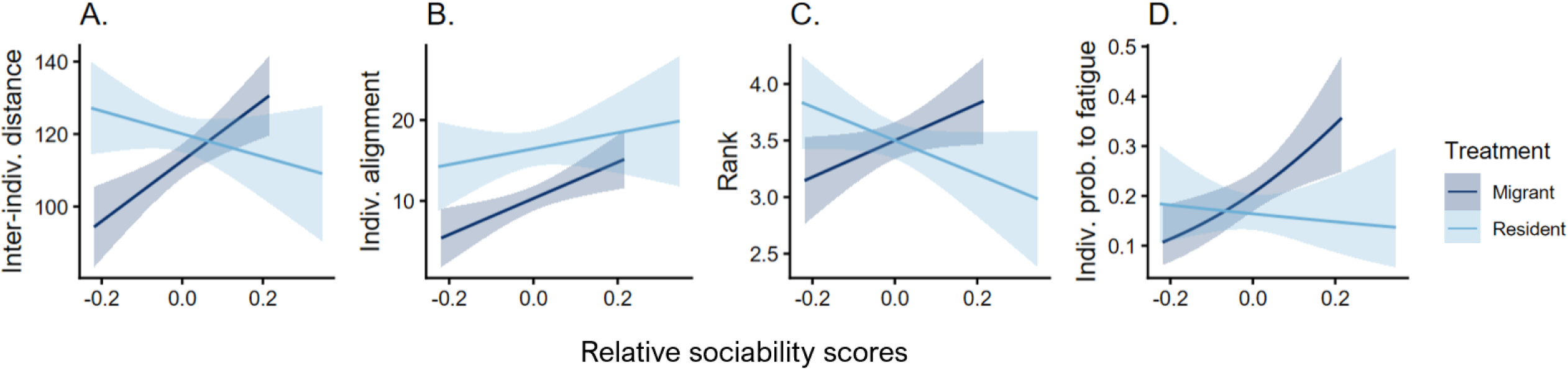
Relationship between individual social tendencies (relative to their group mean) and schooling behaviours in migrant and resident stickleback populations. A. “Inter-individual distance” (mean distance between a focal fish and other members of the school), B. “individual alignment” (deviation in the orientation between a focal fish and the school mean; increasing deviation from 0 indicate decreasing alignment), C. “rank” within the school (1=front of the school, 6= back of the school) and D. “probability to fatigue” (probability that an individual leans at the back of the flow chamber). Regression lines are depicted with their standard errors. Raw data, N=840 observations of 64 individuals (32 migrants and 32 residents).

**Table 1:** Model summary at the individual level examining in pure groups whether variation in individual schooling behaviours (“Inter-individual distance”, “Alignment”, “Rank” within the school and “Probability to fatigue”) was explained by individual phenotype (relative social tendency), population of origin (Migrant or Resident) and their interaction). Estimated effect sizes (β), variances (σ^2^) and (original scale) repeatabilities (R) are reported with their 95% Confidence Intervals (CI). Significant effects are denoted in bold. Trends are inferred from the skewness of CI. Sample size is 840 observations of 64 individuals (32 migrants and 32 residents). Reference category for “Population” is Migrant.

| Schooling trait: | Inter-individual Distance |  | Alignment |  | Rank |  | Prob. to fatigue |  |
| --- | --- | --- | --- | --- | --- | --- | --- | --- |
| Fixed effects | $\beta$ | 95% CI | $\beta$ | 95% CI | $\beta$ | 95% CI | $\beta$ | 95% CI |
| Intercept | **179.81** | **(48.84, 311.49)** | 12.39 | (-22.91, 47.45) | **2.61** | **(0.69, 4.59)** | -9.66 | (-19.12, 1.15) |
| Rel. sociability | **84.96** | **(25.53, 114.13)** | 20.57 | (-2.74, 43.88) | 2.34 | (-0.47, 5.06) | **6.04** | **(0.16, 12.18)** |
| Population | -3.55 | (-37.28, 29.98) | 5.76 | (-3.31, 14.87) | 0.16 | (-0.44, 0.76) | 1.10 | (-1.34, 3.93) |
| Rel. sociability x Population | **-121.60** | **(-206.95, -35.08)** | -7.61 | (-41.29, 25.98) | **-4.45** | **(-8.08, -0.64)** | -7.72 | (-16.08, 0.65) |
| Group mean sociability | -104.24 | (-306.25, 96.67) | -3.34 | (-57.10, 50.81) | 1.40 | (-1.61, 4.31) | 10.86 | (-4.97, 25.94) |
| Random effect | $\sigma^2$ | 95% CI | $\sigma^2$ | 95% CI | $\sigma^2$ | 95% CI | $\sigma^2$ | 95% CI |
| Fish ID | 309.2 | (171.96, 484.37) | 37.76 | (17.12, 61.74) | 0.84 | (0.70, 1.10) | 3.65 | (2.40, 5.29) |
| Group ID | 467.7 | (136.71, 926.33) | 23.98 | (0.00, 56.34) | 0.00 | (0.00, 0.20) | 1.98 | (1.17, 4.71) |
| Trial frame ID | 885.1 | (654.46, 1207.27) | 83.83 | (55.67, 122.62) | 0.00 | (0.00, 0.11) | 1.63 | (1.27, 1.97) |
| Residuals | 1076.6 | (966.69, 1205.54) | 263.61 | (236.95, 295.78) | 2.22 | (1.42, 1.57) | - | - |
| Repeatability (R) | 0.11 | (0.06, 0.17) | 0.03 | (0.00, 0.06) | 0.27 | (0.18, 0.36) | 0.21 | (0.08, 0.31) |

Repeatabilities (adjusted for the fixed effects) showed that *mean inter-individual distance*, *ranking position* and *probability of fatigue* were significantly and moderately repeatable between individuals (range 0.11-0.27), indicating that individuals consistently differ in these schooling traits within- and between trials. In contrast, individual *alignment* was not repeatable over time (Table 1). Note that for *the probability to fatigue,* it did not happen that the all group ended up at the back. Hence, the repeatability of the trait together with the fact that Group ID explained much less variance than Fish ID (1.98 vs. 3.65) indicates that the proportion of fatigued fish is unlikely to result from social attraction at the back.

### Group schooling in pure vs. mixed treatment

Overall, mixed groups appeared to perform slightly worse than pure migrant groups: they had increased inter-individual distances (Fig. 3A) and were less aligned (Fig. 3B) but not different from residents (Table 2). Migrants who are more aligned and compact in pure groups (Fig. 3A-3B, Table 2) tended to be more negatively affected (Fig. 4A-4B, trend in Treatment x Population interaction in Table 3). However, against our initial prediction, migrants were not more likely to take the lead when mixed with residents and fish probability to fatigue was not affected by the treatment (Fig. 4C-4D, no significant Treatment x Population interaction in Table 3).

**Figure 3:**
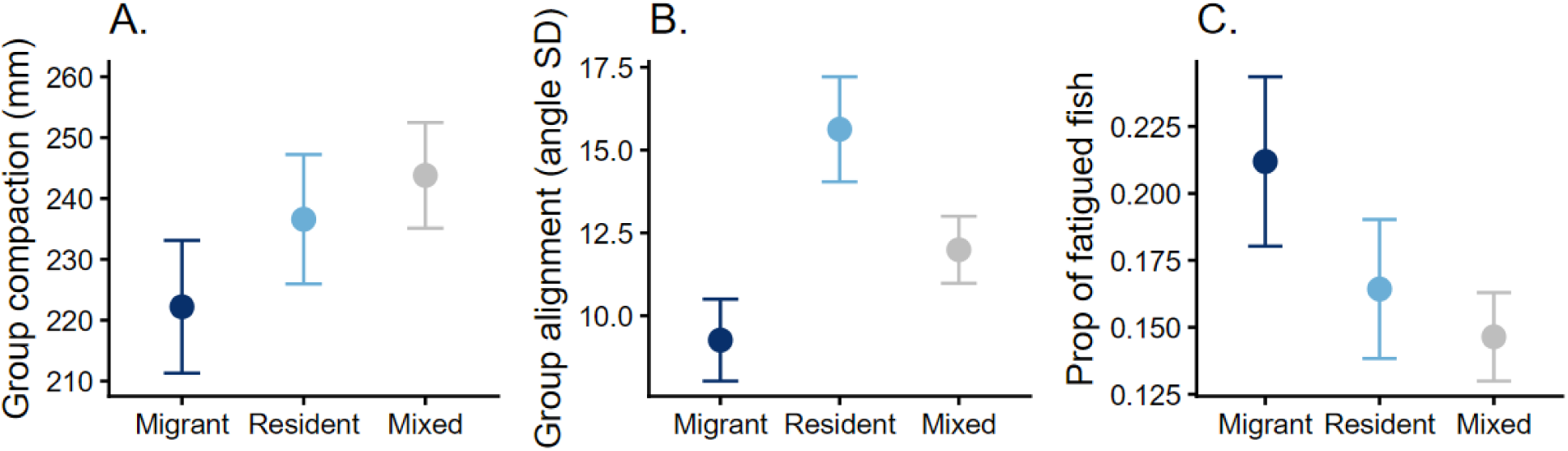
Means and standard errors of schooling behaviours of pure Migrants, pure Residents and Mixed groups. A. “Group compaction” (distance between the first and last fish of the school), B. “Group alignment” (standard deviation of all individual angle deviations from the group mean; increasing values indicate decreasing alignment among school members), and C. “Proportion of fatigued fish” (individuals that leaned at the back of the flow chamber). Raw data, N = 240 observations of 40 schooling trials (10 trials of each pure treatment and 20 trials of mixed treatment).

**Figure 4.**
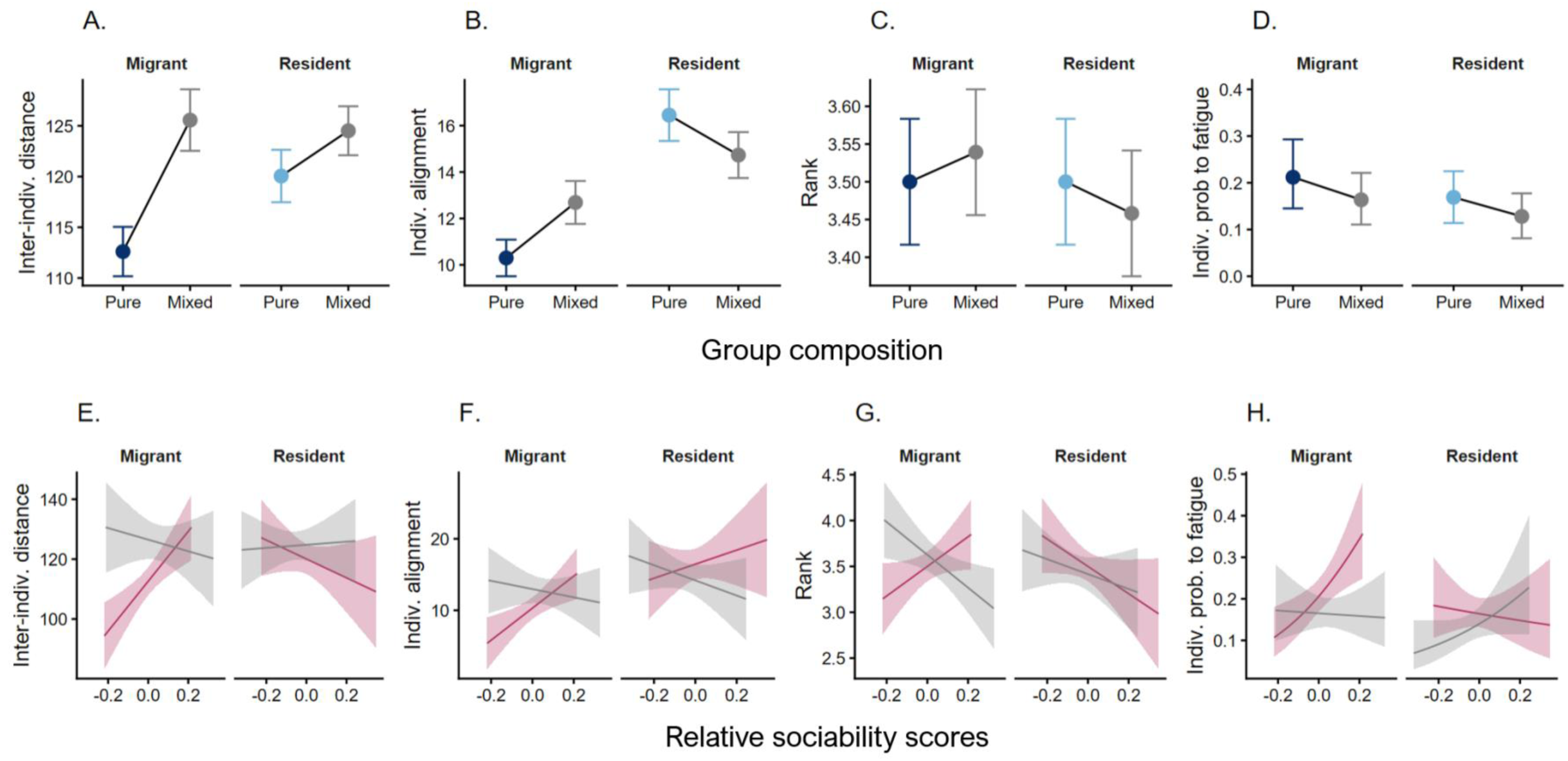
Individual schooling behaviours and its variation across different treatments and sociability tendencies. (A–D) Mean (± SE) schooling behaviours of individual migrant and resident sticklebacks in Pure and Mixed groups, and (E-H) their relationship with individual social tendencies relative to group means. Schooling metrics include: (A and E) *inter-individual distance* (mean distance between a focal fish and other group members), (B and F) *individual alignment* (deviation in orientation between a focal fish and the group mean; higher values indicate lower alignment), (C and G) *ranking position* within the school (1 = front, 6 = back) and (D and H) *probability to fatigue* (probability that an individual leans at the back of the flow chamber) In A-D, colours represent “Pure migrant” (dark blue), “Pure resident” (light blue) and “Mixed” (grey) groups. In E-H, colours represent “Pure” (red) and “Mixed” (grey) groups. Regression lines are shown with their associated standard errors. Raw data, N=1680 observations of 64 individuals (32 migrants, 32 residents) across 20 pure and 20 mixed groups.

**Table 2:** Model summary at the population level examining whether variation in group schooling behaviours (“Group compaction”, “Group alignment” and “Proportion of fatigued fish”) differed between treatments (Migrant, Resident and Mixed groups). Estimated effect sizes (β) and variances (σ^2^) are reported with their 95% Confidence Intervals (CI), with Migrant as the reference ‘Treatment’ group. Significant effects are denoted in bold. Trends are inferred from skewness of CI. Sample size is 240 observations of 40 schooling trials (10 of each pure group and 20 of mixed group).

|  | Group Compaction |  | Group Alignment |  | Prop. Fatigue |  |
| --- | --- | --- | --- | --- | --- | --- |
| Fixed effects | $\beta$ | 95% CI | $\beta$ | 95% CI | $\beta$ | 95% CI |
| Intercept | **222.20** | **(185.35, 259.06)** | **9.26** | **(5.90, 12.57)** | **-1.82** | **-2.74, -0.97)** |
| Treatment: Resident | 14.39 | (-37.73, 66.50) | **6.36** | **(1.61, 11.11)** | -0.23 | (-1.46, 1.02) |
| Treatment: Mixed | 21.27 | (-23.93, 66.47) | 2.73 | (-1.39, 6.84) | -0.33 | (-1.40, 0.75) |
| Random effects | $\sigma^2$ | 95% CI | $\sigma^2$ | 95% CI | $\sigma^2$ | 95% CI |
| Group ID | 2687 | (1253.13, 4467.59) | 11.49 | (0.00, 26.37) | 1.53 | (0.86, 2.90) |
| Residual | 6689 | (5614.20, 8058.28) | 131.35 | (110.40, 157.95) | — | — |

**Table 3:** Model summary examining whether variation in individual schooling behaviours (“mean inter-individual distance”, “alignment”, “ranking position” within the school and “probability to fatigue”) in pure and mixed groups was explained by individual relative sociability, population of origin (Migrant or Resident), the social treatment (Pure vs. Mixed groups) and their interactions. Estimated effect sizes (β) and variances (σ^2^) are reported with their 95% Confidence Intervals (CI). Significant effects are denoted in bold. Trends are inferred from skewness of CI. Raw data, N=1680 observations of 64 individuals (32 M and 32 R) across 20 Pure and 20 Mixed groups. Reference category for “Treatment” is Pure and for “Population” Migrant.

| Schooling traits: | Inter-individual distance |  | Alignment |  | Rank |  | Prob. to fatigue |  |
| --- | --- | --- | --- | --- | --- | --- | --- | --- |
| Fixed effects | $\beta$ | 95% CI | $\beta$ | 95% CI | $\beta$ | 95% CI | $\beta$ | 95% CI |
| Intercept | **181.86** | **(90.79, 273.28)** | 12.02 | (-8.91, 32.83) | **3.76** | **(2.51, 5.01)** | **-4.12** | **(-6.52, -1.82)** |
| Rel. sociability | **81.98** | **(36.49, 127.52)** | **22.13** | **(4.35, 40.02)** | 1.22 | (-0.81, 3.26) | 2.92 | (-0.79, 6.64) |
| Treatment | 9.69 | (-10.00, 29.34) | 2.85 | (-1.73, 7.41) | 0.10 | (-0.14, 0.34) | -0.13 | (-0.56, 0.29) |
| Population | -4.70 | (-32.32, 22.80) | 5.78 | (-0.51, 12.11) | -0.09 | (-0.49, 0.31) | 0.25 | (-0.50, 1.03) |
| Rel. sociability × Treatment | **-100.51** | **(-151.03, -50.07)** | **-26.94** | **(-49.07, -4.82)** | **-3.50** | **(-5.63, -1.37)** | **-5.52** | **(-9.42, -1.74)** |
| Rel. sociability × Population | **-113.94** | **(-178.81, -48.89)** | -10.89 | (-36.76, 14.76) | -2.38 | (-5.24, 0.45) | -1.60 | (-6.69, 3.63) |
| Treatment × Population | 0.37 | (-27.11, 27.95) | -5.52 | (-12.05, 1.03) | -0.09 | (-0.46, 0.28) | -0.19 | (-0.87, 0.48) |

| Rel. sociability × Treatment × Population | **122.37** | **(46.71, 198.16)** | 1.17 | (-31.58, 34.41) | **4.15** | **(1.07, 7.23)** | **9.47** | **(4.09, 15.00)** |
| --- | --- | --- | --- | --- | --- | --- | --- | --- |
| Mean group sociability | -107.33 | (-246.59, 31.42) | -2.62 | (-34.32, 29.26) | -0.36 | (-2.25, 1.53) | 3.95 | (-2.25, 1.53) |
| Random effects | $\sigma^2$ | 95% CI | $\sigma^2$ | 95% CI | $\sigma^2$ | 95% CI | $\sigma^2$ | 95% CI |
| Fish ID | 115.90 | (58.71, 187.77) | 8.95 | (2.35, 17.10) | 0.95 | (0.74, 1.29) | 0.28 | (0.40, 0.65) |
| Group ID | 437.10 | (185.80, 714.71) | 12.28 | (0.00, 26.78) | — | — | — | — |
| Trial frame ID | 979.30 | (785.13, 1226.17) | 84.16 | (62.92, 111.08) | — | — | — | — |
| Residuals | 1409.90 | (1305.75, 1519.77) | 284.00 | (263.09, 309.14) | — | — | 2.64 | (1.57, 1.68) |
| Repeatability (R) | 0.04 | (0.02, 0.06) | 0.02 | (0.00, 0.04) | 0.10 | (0.06, 0.14) | 0.07 | (0.00, 0.00) |

### Personality-dependent plasticity in schooling in pure vs mixed groups

Our analyses further showed that sociability and plastic adjustments to changes in the group social composition were not linked: In mixed groups, sociability did not covary with schooling behaviour. This is particularly pronounced in migrants, for which the previous relationship found between relative sociability and schooling behaviours collapsed when they were mixed with residents (Fig. 4E-H, and three-way interactions in Table 3). These patterns were confirmed when populations were analysed separately (not shown).

Across treatments, repeatabilities (adjusted for the fixed effects) of schooling behaviours dropped considerably, indicating that individuals were not able to maintain their average behaviour when mixed with other ecotypes (Table 3).

## Discussion

This study demonstrated that schooling behaviour is shaped by variation at multiple levels from individual personality to group composition to population. At the population level, migrants exhibited higher alignment and tended to form more compact schools than residents, but fatigued more. When mixed, group schooling performance decreases with mixed groups tending to have worse compaction and intermediate alignment but were not significantly different from resident groups. Analyses at the individual level, uncovered that this effect was due to overall declining schooling performance of migrants, potentially due to the disruption of phenotype-dependent schooling in mixed groups. Although the size of the testing area was relatively small and may have somewhat constrained schooling behaviour, the setup still uncovered significant variation in school compaction and individual probability to fatigue. We discuss the implications of our findings and the importance of studying behavioural variation across multiple hierarchical levels to better understand the mechanisms underlying collective behaviour.

### Population-Dependent Schooling Behaviour

Previous studies have shown that freshwater sticklebacks are less capable of performing schooling behaviour than anadromous sticklebacks (Wark et al. 2011, Greenwood et al. 2013, 2016, Di-Poi et al. 2014). The reduced tendency to school in freshwater sticklebacks is argued to reflect an adaptation to complex habitats containing many shelters, where fish may not need to aggregate to find safety from predators (Wark et al. 2011). Although our study can only compare one pair of populations (anadromous vs. pooled resident populations), which limits its generalisability, our results are in line with these previous findings: fish within resident schools were less capable of schooling (they were less aligned and less cohesive) than migrant groups. This confirms that 50-60 years of isolation are sufficient to induce rapid behavioural divergence between Dutch landlocked resident and anadromous populations, with residents on their way to losing migration-related behaviours (Ramesh et al. 2021, 2022). Given that schooling behaviour is known to have a genetic basis in sticklebacks, with the *Ectodysplasin A* (Eda) gene controlling the main phenotypic attributes of freshwater sticklebacks (such as reduced lateral plating and reduced schooling) (Colosimo et al. 2005, Jones et al. 2012, Greenwood et al. 2013, 2016), and that we used F1 fish raised under similar environmental conditions, the observed differences in schooling must be underpinned by genetic differentiation.

Mixed groups performed worse overall: they were less aligned than migrants but were not different from residents and had lower compaction, which was not significantly different from the pure groups. Contrary to our expectations, migrants were not more likely to take leading positions in mixed groups. Interestingly, mixed groups had lower probability to fatigue. This may be caused by habituation as fish were tested on average four times and mixed groups were always tested after the pure groups (Supplementary Material S1).

All together, these results indicate a lack of adaptive plasticity or social modulation by residents or migrants in mixed groups. We previously found that residents and migrants did not adjust their large-scale movement tendencies to group composition (Gismann et al. 2024). One likely explanation is that mixing populations increases within-group heterogeneity, disrupting coordination. Although these F1 fish did not differ in size, residents may lack the physical capacities to coordinate movements with migrants (which could result from their genetic differentiation), leading to reduced group functioning. This aligns with modelling studies showing that increased behavioural variation can hinder collective motion (Del Mar Delgado et al. 2018). Furthermore, naturally formed schools are typically characterized by high uniformity in terms of size or physiological status (Kasumyan and Pavlov 2023c), including sticklebacks (e.g., Ranta et al. 1992). This uniformity is achieved by the passive exit from the schools of individuals with reduced swimming capacities (Kasumyan and Pavlov 2023c). Consequently, it seems very unlikely that recently diverged residents and anadromous individuals will form cohesive schools once river connectivity will be restored. These differences may ultimately favour the emergence of partial migration in our system.

### Personality-dependent schooling behaviour

At the individual level, sociability strongly covaried with schooling in migrants. As predicted, less social fish were more often in the leading position and exhibited better alignment. However, unexpectedly, these asocial fish swam closer to each other and fatigued less, indicating overall better swimming and schooling abilities than more social fish. For pelagic species that need to undergo migration in open habitats, leading positions, which experience more resistance from the environment, may thus be occupied by individuals that engage more in solitary swimming (i.e., less social individuals) and have higher swimming capacities. Conversely, more social fish who appeared here to be weaker swimmers stayed more at the back of the school. This is similar to a peloton of cyclists, where weaker riders cycle behind leaders to expend less energy and/or be able to maintain the speed of stronger riders (Trenchard 2011). In support, a previous study showed that more asocial sticklebacks swam faster, were more aligned, and therefore ended up more often in front (Jolles et al. 2017).

Our analyses further reveal that all schooling traits, except alignment, were significantly repeatable among individuals tested in pure groups in a range that is expected for behavioural traits (range 0.11-0.27) (Bell et al. 2009). This repeatability likely reflects stable social dynamics shaped by consistent differences in sociability, especially in migrant groups. Individual differences in social responsiveness can promote the emergence of structured leader–follower dynamics and, as previously shown in sticklebacks, this dynamics is further reinforced by positive feedback loops, where more responsive social/shyer individuals make asocial/bolder individuals to lead even more (Harcourt et al. 2009). However, in mixed groups, the relationship between sociability and schooling behaviour disappeared and repeatability of schooling behaviours decreased across contexts. Mixing populations may thus hamper social signalling, disrupt positive social feedback and reduce the propensity of asocial migrants to initiate leadership.

### Variation across hierarchical levels

Research on animal behaviour increasingly emphasises that variation is structured at multiple, nested hierarchical levels, each with distinct biological meaning (Dingemanse et al. 2010, Hertel et al. 2020). In this study, we show that schooling in sticklebacks is shaped by variation at the among-population and among-individual within-population levels. In homogeneous anadromous groups, individual differences in sociability drive self-organization group structure and overall schooling cohesion in migrants. By introducing heterogeneity through mixed ecotypes, we found that structure of phenotype-dependent schooling is disrupted, leading to reduced group schooling ability. This indicates that collective behaviour is constrained by the least performing individuals within a group, rather than driven by its highest-performing individuals.

## Conclusions

Our findings indicate that schooling tendencies in three-spined sticklebacks are strongly shaped by the structure of phenotype-dependent schooling in which individual differences in sociability and potentially other behavioural traits interact to organise group collective movement. This structured organisation is absent in residents, which exhibit lower schooling ability and reduced sociability, potentially underlined by genetic divergence. Our study thus adds to the growing literature that considering behavioural variation across multiple hierarchical levels is an important future avenue to understand complex social systems.

## Acknowledgements

We thank Rik Nienhuis for help with the experiment, data collection and analysing the first part of the videos and Jasper Wietses for analysing the second part of the videos. We thank Dennis de Worst and Willem Diederik for help with fish care and advice on experimental design. We thank Mariana M. Domingues for help with the personality tests. We thank Peter Paul Schollema, at the Water Authorities Hunze en Aa’s and Jeroen Huisman at van Hall Larenstein, University of Applied Sciences, for help with acquiring wild sticklebacks. We also thank two anonymous reviewers who helped improve the manuscript.

## Funding

This work is supported by PhD fellowship of the Adaptive Life program of the University of Groningen to AR, by funding from SNSF grant 310030_207448 to AR and B Taborsky and from the Netherlands Organization for Scientific Research to FJ Weissing and MN (NWO-ALW; ALWOP.668). This work was also supported by grants from the Gratama Foundation to AR (2020GR040), the Dr. J.L. Dobberke Foundation (KNAWWF/3391/1911), and the Waddenfonds - Ruim Baan voor Vissen 2 (01755849/WF-2019/200914).

## Ethics approval

Sampling of wild animals and handling methods were done following a fishing permit from Rijksdienst voor Ondernemend Nederland (the Netherlands) and an angling permit from the Hengelsportfederatie Groningen-Drenthe. Animal housing and behavioural tests adhered to the project permit from the Centrale Commissie Dierproeven (the Netherlands) under the license number AVD1050020174084. All methods were carried out under the applicable international, national, and institutional guidelines for the use of animals (Art. 9, Wet op de Dierproeven & European directive 2010/63/EU).

## Data availability

Analyses reported in this article can be reproduced using the data provided by https://figshare.com/s/d33ea8301f447ac97ccc

## Notes

https://figshare.com/s/d33ea8301f447ac97ccc

### Competing Interest Statement

The authors have declared no competing interest.

