## Supplementary Material S1 for "Phenotype-dependent schooling behaviour across hierarchical levels in three-spined sticklebacks"

**Supplementary material S1 – Probability to fatigue as a function of number test repeats**

Table S1 and Figure S1 show that fatigue probability decreased significantly across repeats, indicating that individuals were less likely to fatigue in later repeats. Pure trials were always performed first and were within 1-3 repeats and later repeats comprised mostly of mixed group trials.

**Table S1:** model summary examining Probability to fatigue as a function of the number of test repeats. A generalized linear mixed-effects model with a binomial error structure showed a negative effect of repeat number (1 to 6) on the probability to fatigue. Estimates (β) and variance (σ^2^) effect are given with their 95% confidence interval (CI). Significant values are depicted in bold.

|  | **Probability of fatigue** | |
| --- | --- | --- |
| Fixed effect | β | 95% CI |
| Intercept | **-1.283** | **(-1.687, -0.900)** |
| repeat | **-0.227** | **(-0.357, -0.099)** |
| Random effect | σ^2^ | 95% CI |
| Fish ID | 0.794 | (0.458, 1.372) |

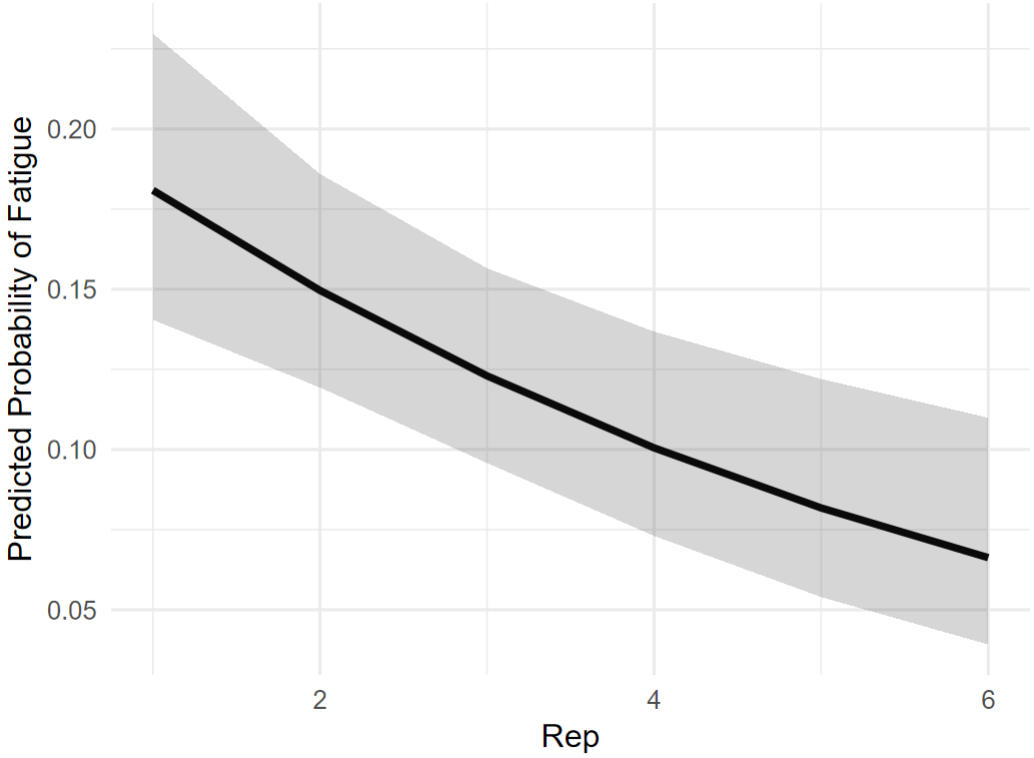

**Figure S1**: Predicted probability to fatigue as a function of number of repeats per fish. Shaded areas represent 95% confidence intervals around model predictions.
